# Condition-matched in silico prediction of drug transcriptional responses enables mechanism-guided screening and combination discovery

**DOI:** 10.64898/2026.03.27.714886

**Authors:** Meisheng Xiao, Yiping He, Jianhua Hu, Fei Zou, Baiming Zou

**Author notes:** Contributing authors.

## Abstract

Perturbational transcriptomics links therapeutic compounds to cellular mechanisms and provides a powerful framework for drug discovery, but experimentally profiling transcriptional responses across diverse cell states, doses and durations is costly and often infeasible. Here we present DEPICT (Drug rEsponse Prediction in transCriptomics with Transformers), a deep learning framework that predicts condition-matched drug-induced transcriptional responses from baseline gene expression, perturbation settings and complementary drug representations. Using the LINCS L1000 dataset, DEPICT generalized to unseen drugs and cell types and outperformed five baseline strategies and two recent deep learning models. In the most challenging unseen-cell evaluation, DEPICT was the only model to surpass all baselines, improving differential-expression prediction accuracy and reducing perturbed-expression prediction error by 30.3% and 36.8%, respectively, relative to the next-best deep model. In a non-small cell lung cancer (NSCLC) case study, DEPICT-enabled virtual screening prioritized compounds predicted to reverse disease-associated transcriptional signatures. Notably, 13 of the top 20 prioritized compounds had either previously entered NSCLC-related clinical trials or been validated in NSCLC studies, supporting the translational relevance of the predicted perturbational profiles. DEPICT further enabled condition-matched drug synergy prediction and mechanistic exploration when experimentally matched profiles were unavailable. Together, these results show that accurate, condition-matched in silico perturbation profiling can scale transcriptomics-driven hypothesis generation for drug repurposing and combination discovery.

## Introduction

Precision oncology seeks to identify therapies tailored to the molecular state of an individual tumor [1]. In this context, identifying therapies capable of reversing tumor-specific transcriptional programs has emerged as a promising strategy for guiding treatment selection and improving clinical outcomes. One powerful approach to enable such strategies is perturbational transcriptomics, which compares a baseline cellular state with the transcriptional state induced after treatment at the systems-biology level [2–4]. A central but often underappreciated feature of perturbational data is that a transcriptional signature is not a fixed property of a compound. Rather, the perturbational response can vary substantially with cellular context and exposure conditions, including dose and duration [5, 6]. This context dependency is particularly pronounced in cancer, where heterogeneous and evolving tumor states can alter therapeutic sensitivity and resistance [7, 8]. Accordingly, identifying effective therapies requires perturbational profiles measured under matched biological and pharmacological conditions. When only mismatched reference signatures are available, critical drug effects may be obscured, thereby limiting the translation of preclinical hypotheses into successful therapeutic outcomes [9, 10]. Thus, the biologically meaningful unit for preclinical discovery is not simply a drug-induced signature, but a condition-matched drug response signature. However, the combinatorial space defined by biological context, compound, dose, and duration far exceeds what can be experimentally profiled. As a result, many clinically relevant conditions remain unmeasured, creating a fundamental bottleneck in leveraging perturbational transcriptomics for translational research and motivating the need for accurate in silico prediction of condition-matched transcriptional responses.

To address this challenge, a growing number of predictive models have been developed by leveraging large scale perturbational resources such as LINCS L1000 [11], chemical databases such as PubChem [12], and recent advances in machine learning [13, 14]. These approaches aim to predict post-perturbation transcriptional states from baseline gene expression and drug features [15–20]. However, their translational utility remains limited. Many models rely on single-source drug representations [21], do not explicitly account for exposure conditions such as dose and duration [15], or generalize poorly to previously unseen biological contexts [22, 23]. Consequently, predicted transcriptional responses may fail to capture the context-specific perturbational effects required to guide preclinical decision-making and therapeutic prioritization.

To overcome these limitations, we developed DEPICT (Drug rEsponse Prediction in transCriptomics with Transformers), a deep learning framework for predicting drug-induced transcriptional responses under specified biological and pharmacological conditions. By integrating baseline gene expression, perturbation settings, and complementary representations of drug structure and biomedical knowledge, DEPICT generates condition-matched perturbational profiles that can support downstream applications including therapeutic prioritization and drug combination discovery. We evaluated DEPICT using large scale perturbational data from LINCS L1000 [11] and demonstrated its ability to predict transcriptional responses for previously unseen drugs and cellular contexts. We further show that DEPICT enables mechanism-guided therapeutic prioritization in non-small cell lung cancer (NSCLC), supports condition-matched drug synergy prediction when experimentally matched profiles are unavailable, and reveals transcriptional landscapes that may guide hypothesis generation for combination therapies. Together, these findings suggest that condition-matched in silico perturbation profiling can provide a practical path toward scaling transcriptomics-driven hypothesis generation for preclinical therapeutic discovery.

## Results

DEPICT is a transformer-based deep learning framework that aims to accurately predict transcriptional responses induced by drugs. We trained, validated, and tested DEPICT on preprocessed level-3 LINCS L1000 (GSE92742) perturbation data [11]. This dataset contains 836,649 perturbation profiles and 46,428 baseline profiles from 2,875 plates. It covers 82 cell lines and 17,203 drugs across various levels of dose and duration. To provide a baseline expression for every perturbation, we paired each perturbation profile with a randomly selected baseline profile from the same plate. DEPICT takes the baseline expression of the 978 landmark genes, a dual-view drug representation—capturing local chemical substructures via Morgan fingerprints and broad biomedical properties via large language model (LLM) embeddings—along with the dose and duration of the perturbation as inputs to predict the expression of those 978 genes after perturbation.

By generating perturbation profiles under matched conditions, DEPICT supports preclinical analyses that rely on changes in transcription. In this study, we demonstrated the superior predictive accuracy of DEPICT through extensive benchmarking against baselines and deep learning frameworks, and we then applied it to translational downstream tasks, including screening for candidates that reverse the NSCLC disease signature, predicting synergy for drug doublets, and conducting exploratory analyses based on the predictions. In oncology, these capabilities are directly relevant to prioritizing therapies for molecularly defined tumors, anticipating pathway-level effects linked to response and resistance and narrowing the search space for combinations that are most likely to be synergistic in a given tumor context. Additionally, the architectural design choices in DEPICT enable other meaningful preclinical applications. For example, explicitly modeling dose and duration supports future studies regarding the relationships between dose and response or the scheduling of treatments, while modeling at the individual gene level provides a natural handle to interpret features and attribute biological meaning. An overview of the DEPICT workflow is illustrated in Fig. 1.

**Fig. 1:**
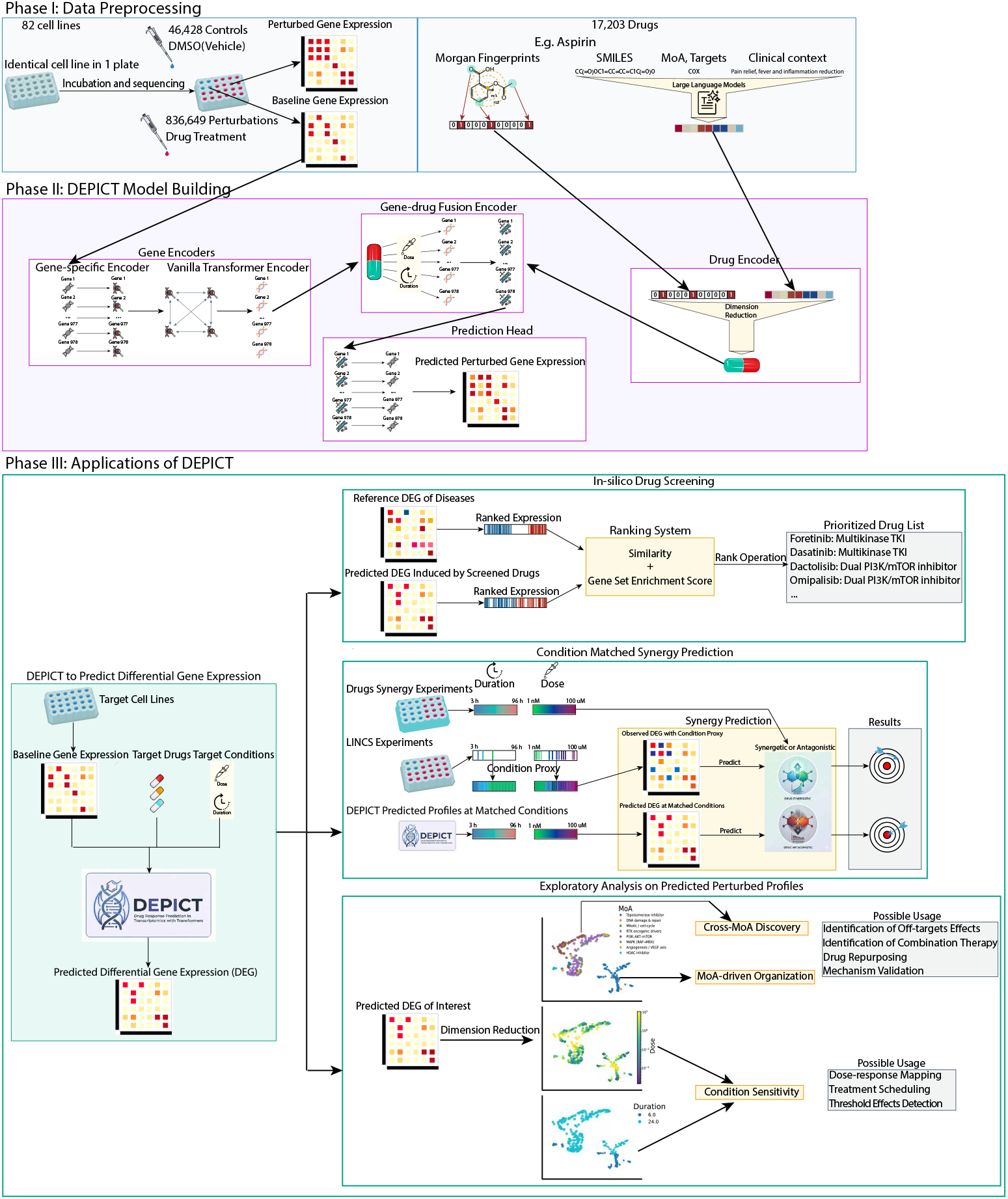
Overview of DEPICT

### Model performance comparison

We benchmarked DEPICT across three distinct data splitting strategies, each representing a specific translational challenge. The random split evaluates the model’s ability to impute untested experimental condition, such as missing doses or durations, for known drug-cell pairs. The drug split represents the challenge of predicting the effects of novel, unprofiled compounds. Finally, the cell split measures the capacity to forecast responses in entirely uncharacterized biological contexts, such as newly derived cell lines or rare cancer lineages.

We summarized the comprehensive performance of all models in Table 1. We evaluated predictive accuracy using Mean Squared Error (MSE), Pearson Correlation Coefficient (PCC), and Coefficient of Determination (*R*^2^). In our notation, Δ denotes metrics calculated on the differential expression relative to the baseline state. To establish a rigorous performance floor, we compared DEPICT with simple evaluation methods, such as using baselines as predictions, alongside recent deep learning frameworks, including TranSiGen [15] and PRnet [16]. (See Methods for detailed descriptions of all simple evaluation methods).

**Table 1:** Model performance across three generalization schemes.

| Model | MSE | $R^2$ | PCC | $\Delta R^2$ | $\Delta PCC$ |
| --- | --- | --- | --- | --- | --- |
| **(a) Cell split** |  |  |  |  |  |
| Naive | 1.462 (0.103) | 0.766 (0.010) | 0.884 (0.005) | — | — |
| TM | 2.595 (0.342) | 0.574 (0.044) | 0.760 (0.029) | -1.457 (0.304) | 0.338 (0.019) |
| TM-D | 2.452 (0.238) | 0.599 (0.027) | 0.776 (0.018) | -1.326 (0.229) | 0.360 (0.014) |
| PRnet | 3.014 (0.394) | 0.503 (0.064) | 0.778 (0.009) | -1.762 (0.462) | 0.296 (0.019) |
| TranSiGen | 1.570 (0.177) | 0.746 (0.026) | 0.864 (0.015) | -0.371 (0.204) | 0.435 (0.029) |
| DEPICT | 0.992 (0.090) | 0.844 (0.008) | 0.918 (0.004) | 0.283 (0.025) | 0.567 (0.006) |
| **(b) Drug split** |  |  |  |  |  |
| Naive | 1.349 (0.021) | 0.778 (0.004) | 0.890 (0.002) | — | — |
| TM | 2.265 (0.014) | 0.620 (0.003) | 0.789 (0.002) | -1.333 (0.033) | 0.348 (0.002) |
| TM-C | 1.409 (0.026) | 0.770 (0.004) | 0.878 (0.003) | -0.286 (0.018) | 0.461 (0.003) |
| PRnet | 1.839 (0.028) | 0.696 (0.005) | 0.849 (0.002) | -0.608 (0.024) | 0.377 (0.003) |
| TranSiGen | 0.799 (0.016) | 0.871 (0.003) | 0.932 (0.002) | 0.381 (0.003) | 0.630 (0.002) |
| DEPICT | 0.801 (0.011) | 0.871 (0.002) | 0.933 (0.003) | 0.384 (0.002) | 0.623 (0.002) |
| **(c) Random split** |  |  |  |  |  |
| Naive | 1.373 (0.004) | 0.772 (0.001) | 0.888 (0.001) | — | — |
| TM | 2.270 (0.003) | 0.618 (0.001) | 0.788 (0.001) | -1.296 (0.005) | 0.350 (0.001) |
| TM-CD | 1.384 (0.006) | 0.776 (0.001) | 0.884 (0.001) | -0.223 (0.003) | 0.501 (0.001) |
| PRnet | 1.841 (0.007) | 0.695 (0.001) | 0.848 (0.001) | -0.579 (0.009) | 0.382 (0.001) |
| TranSiGen | 0.775 (0.003) | 0.874 (0.001) | 0.934 (0.001) | 0.401 (0.002) | 0.645 (0.001) |
| DEPICT | 0.779 (0.004) | 0.874 (0.001) | 0.934 (0.001) | 0.407 (0.002) | 0.640 (0.001) |

Values are mean (SD) over 5 repetitions.
Naive = Baselines as Predictions; TM = Train set Mean; TM-C = Train set Mean across Cell; TM-D = Train set Mean across Drug; TM-CD = Train set Mean across Cell-Drug Pair.

As shown by the naive baseline in Table 1, the average expression difference between the baseline and perturbed states is modest. This indicates that a cell’s baseline expression inherently dictates a large portion of its post-perturbation profile. Furthermore, the overall results identify the unseen cell split as the most challenging generalization scenario. When assessing generalizability to these unseen cell lines (Table 1a), DEPICT outperformed all other models across all metrics and emerged as the only machine learning framework to exceed all baseline evaluations. Conversely, competing deep learning models underperformed the naive baseline, which simply outputs the baseline profile as the final prediction. Bridging this generalization gap is clinically important, as an unseen cell in a translational pipeline serves as a direct proxy for encountering a novel biological environment, such as a new patient context, a rare cancer lineage, or an emerging disease model.

Within the unseen cell evaluation, DEPICT achieved a 32.1% reduction in MSE and a 10.2% improvement in *R*^2^ compared to the naive baseline. When evaluated against the next best deep learning framework, DEPICT reduced the MSE by 36.8% and increased ΔPCC by 30.3%, indicating superior accuracy in predicting both the magnitude and directionality of expression signals induced by perturbations. Crucially, DEPICT was the sole framework to obtain a positive Δ*R*^2^. This metric confirms that DEPICT predicts the drug-induced differential gene expression (DGE), whereas competing models fail to distinguish the perturbation effect from the inherent baseline expression. Beyond unseen cells, DEPICT also achieved top-tier predictive accuracy in drug and random splits. This consistent performance across all evaluation scenarios establishes DEPICT as a generalizable framework capable of predicting drug responses for both novel compounds and new cell lines under required perturbational conditions.

### Mechanism-guided virtual screening identifies candidates that reverse NSCLC transcriptional signatures

Lung cancer remains the most lethal malignancy worldwide, with non-small cell lung cancer (NSCLC) accounting for approximately 84% of all diagnoses [24, 25]. The biological diversity and rapid emergence of resistance in NSCLC present significant hurdles for therapy development [26, 27]. To evaluate whether condition-matched in silico profiles support actionable hypothesis generation, we tested DEPICT in a large scale, NSCLC-focused virtual screening setting. Specifically, we prioritized 17,203 compounds based on their predicted ability to reverse a 690-gene NSCLC disease signature in the A549 cell line (see Methods). We defined successful screening by two criteria: (i) the mechanistic concordance of top-ranked candidates with established NSCLC-relevant targets and pathways, and (ii) the preferential ranking of clinically investigated NSCLC therapies. Here, we report mechanistic concordance analyses and quantitative retrieval metrics, highlighting representative top-ranked therapeutic candidates (Fig. 2 and Table 2).

**Table 2:** Validated candidate compounds for NSCLC from the top-20 list.

| Drug | MoA | Evidence |
| --- | --- | --- |
| MK-2206 | Akt inhibitor | Phase II [33] |
| Dasatinib | Multikinase TKI | Phase II [34] |
| AZD-8330 | MEK inhibitor | Phase I [35] |
| PHA-793887 | CDK inhibitor | Phase I [36] |
| ZSTK-474 | Pan PI3K inhibitor | Phase I [37] |
| Foretinib | Multikinase TKI | Phase I [38] |
| NVP-BEZ235/dactolisib | Dual PI3K/mTOR inhibitor | Phase I [39] |
| GSK-2126458/omipalisib | Dual PI3K/mTOR inhibitor | Phase I [40] |
| OSI-027 | mTOR inhibitor | Preclinical [41] |
| PI-103 | PI3K/Akt/mTOR inhibitor | Preclinical [42] |
| Torin-2 | mTOR inhibitor | Preclinical [43] |
| CD-437 | Apoptosis | Preclinical [44] |
| Midostaurin | Multikinase TKI | Preclinical [45] |

**Fig. 2:**
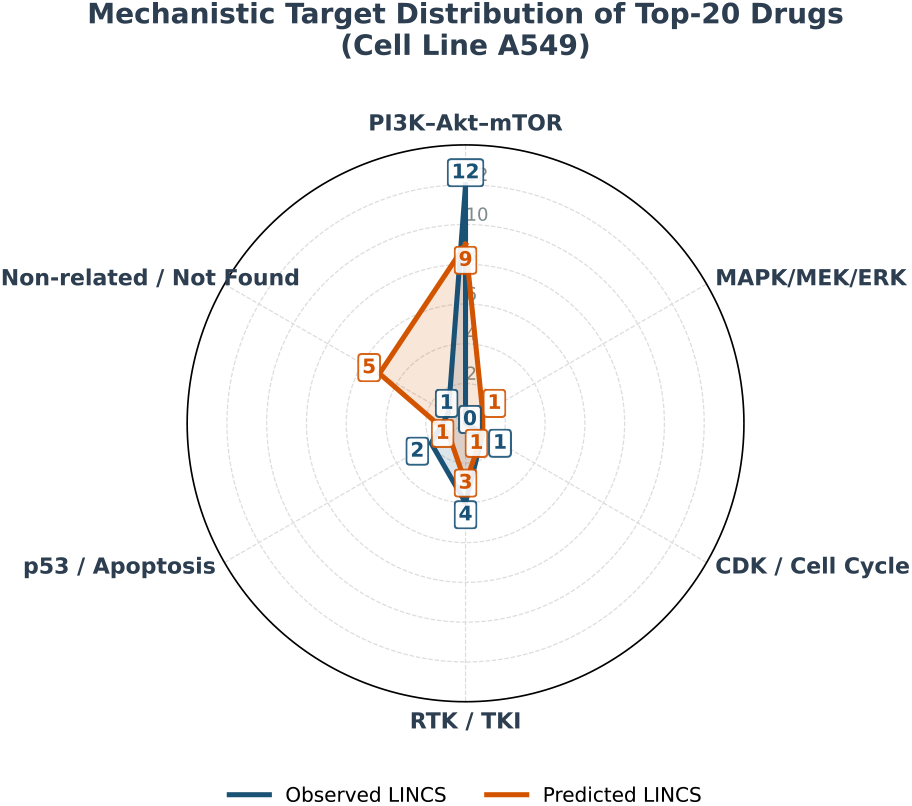
Drug MoA radar plot

As shown in Fig. 2, 15 of the top 20 drugs prioritized using the predicted LINCS L1000 profiles exhibit mechanisms of action (MoAs) relevant to NSCLC treatment. Nine of these drugs inhibit the PI3K–Akt–mTOR pathway, one of the most important signaling axis implicated in lung cancer progression [28]. To validate the observations made by predictions, we repeated the same analysis using observed LINCS L1000 data.

19 of the top 20 drugs have relevant MoAs, including 12 PI3K–Akt–mTOR inhibitors. Both predicted and observed results reveal a similar mechanistic pattern. Most top-ranked compounds are functionally associated with NSCLC, with strong enrichment for PI3K–Akt–mTOR pathway inhibitors. The enrichment for PI3K–Akt–mTOR inhibitors aligns with known biology. First, A549 is a KRAS-mutant, LKB1-deficient NSCLC line that exhibits marked sensitivity to PI3K–Akt–mTOR pathway inhibition [29, 30]. Second, the PI3K–Akt–mTOR pathway serves as a major convergence node for multiple upstream oncogenic signals, and its activation influences the expression of many other genes [31]. Third, PI3K–Akt–mTOR related perturbations are well reflected in the LINCS L1000 dataset [32]. One notable difference in patterns is that the prediction-based ranking includes five compounds with unknown or unrelated MoAs in the context of NSCLC. For example, the cardiovascular metabolite MRE-269 emerges as a potential drug repurposing candidate, whereas JWE-035 and VU-0418947-2 highlight preclinical oncology pathways (Aurora kinase and hypoxia/HIF) not currently utilized in NSCLC care. Finally, JW-7-24-1 and the probe BRD-K26381032 lack any annotated targets. Because their strong prioritization is driven by their predicted capacity to invert the transcriptomic disease signature, these diverse molecules serve as highly specific, mechanism-driven leads for subsequent in vitro validation. In addition to comparing MoA patterns, we evaluated the overlap between predicted and observed drug rankings across the 17,203 compounds. The overlap of the top-ranking lists included two drugs (20%) in the top 10, five (25%) in the top 20, and sixteen (32%) in the top 50. The above consistencies between predicted and observed rankings demonstrate that DEPICT can effectively guide preclinical drug screening, especially when needed experimental conditions are unavailable in existing datasets. The detailed list of top-20 prioritized candidates can be found in Appendix Table B5.

Table 2 summarizes 13 compounds from the top-20 prioritized list with prior evidence related to NSCLC, including 8 that had advanced to clinical trials and 5 with preclinical support. The clinical-stage compounds (MK-2206, dasatinib, AZD-8330, PHA-793887, ZSTK-474, foretinib, dactolisib, and omipalisib) indicate that DEPICT recovers candidates that have already been considered relevant in lung cancer drug development. The additional hits with preclinical evidence (OSI-027, PI-103, torin-2, CD-437, and midostaurin) extend this pattern beyond compounds that entered trials and suggest that DEPICT also prioritizes biologically plausible candidates linked to NSCLC-relevant pathways. Among the remaining prioritized compounds, PI-828 and KU-0060648 had limited direct NSCLC evidence in our review and are therefore better viewed as exploratory leads for follow-up validation [46, 47].

These findings support the use of predicted condition-matched perturbation profiles as a pre-screening layer that narrows a large candidate space to a smaller set of translationally relevant and hypothesis-generating therapies. DEPICT may also be useful for screening novel compounds that have not yet been experimentally profiled. When integrated with de novo drug design pipelines, it could help prioritize candidate therapies for specific diseases at scale and thereby support early stage therapeutic discovery. Overall, DEPICT-generated LINCS L1000 predictions illustrate the potential of large scale, condition-specific in silico drug screening for drug repurposing and early stage candidate prioritization.

### Synergy prediction using condition-matched predicted profiles

Drug combinations are a mainstay of modern oncology, essential for overcoming resistance and enhancing therapeutic efficacy [48–51]. Because the vast combinatorial space of potential candidates makes exhaustive experimental screening expensive, in silico synergy prediction has emerged as a practical tool to narrow the search to the most promising regimens [52, 53]. However, developing accurate predictive models is hindered by a data mismatch problem. Experimental synergy labels are rigorously curated across complex, multi-dose and multi-duration settings. In contrast, high-throughput transcriptomic datasets, such as LINCS L1000, are highly sparse and rarely contain these exact perturbation conditions. Using these mismatched transcriptomes introduces measurement error that compromises predictive accuracy. DEPICT directly resolves this by generating condition-matched profiles, providing more reliable inputs for robust synergy prediction.

In this analysis, we used both predicted and observed LINCS L1000 transcriptional profiles as input features to predict whether drug pairs are synergistic or antagonistic in the HT29 cell line. The drug synergy reference dataset [54] was curated under doses and durations that differ from those in the LINCS framework. Therefore, for the observed LINCS data, we used perturbations with the closest available experimental conditions as substitutes when exact matches were not available. In contrast, for the DEPICT-predicted LINCS data, we directly generated profiles under the same experimental conditions as in the reference dataset. Ridge logistic regression (RLR) and random forest (RF) were then used to classify the drug synergy using either the observed or the predicted LINCS profiles. Model performance was evaluated using leave-one-out strategy, with AUC (Area Under the Curve), PR-AUC (Area Under the Precision-Recall Curve), Accuracy (ACC) and macro F1 score, as the performance evaluation metrics.

As shown in Table 3, classifiers trained on predicted transcriptional responses consistently and remarkably outperform those using observed LINCS data under closest condition proxy across all evaluation metrics. This performance gap can be attributed to the absence of exact condition-matched perturbation profiles in the observed LINCS data, as the closest available observed profiles may approximate expression values but can distort the true direction or strength of perturbational signals. These results indicate that DEPICT effectively captures the influence of experimental conditions and that dose and duration are critical factors for accurate drug synergy prediction. Overall, the ability of DEPICT to generate condition-matched transcriptional profiles offers a more flexible and accurate alternative to experimentally observed data when exact matched conditions are unavailable.

**Table 3:** Performance comparison between using predicted and observed transcriptional responses for synergy predictions.

| Model | Data Source | AUC | PR-AUC | ACC | Macro F1 |
| --- | --- | --- | --- | --- | --- |
| RLR | Predicted LINCS | 0.844 | 0.714 | 0.802 | 0.784 |
|  | Observed LINCS | 0.786 | 0.687 | 0.752 | 0.717 |
| RF | Predicted LINCS | 0.855 | 0.770 | 0.802 | 0.782 |
|  | Observed LINCS | 0.777 | 0.639 | 0.745 | 0.714 |

RLR = Ridge Logistic Regression RF = Random Forest

Furthermore, the reported results from DeepDDS [55], a machine learning framework that predicts drug pair synergy using baseline gene expression from CCLE [56] and graph-based drug embeddings, support the reliability of our findings. The random forest performance reported by DeepDDS (AUC = 0.82; PR-AUC = 0.81; ACC = 0.74) is comparable to our classifiers trained on predicted LINCS profiles, underlying both the validity of DEPICT’s performance and the potential of using transcriptional perturbation data to enhance drug synergy prediction.

### Exploratory analysis on predicted perturbations

To investigate how the model organizes the complex landscape of drug-induced changes, we used 2D UMAP embeddings of DEPICT-predicted differential gene expression (DGE) to explore how lung-cancer relevant drugs are organized. The analysis included 1,110 perturbations on the NSCLC cell line A549, from 166 unique drugs with mechanisms associated with lung cancer.

In the UMAP plots of 166 compounds (Fig. 3A), perturbations were organized into four primary, well-separated areas, each characterized by distinct structural patterns that underscore the framework’s ability to prioritize different biological and experimental signals. Two of these areas correspond to HDAC inhibitors (Area II) and topoisomerase inhibitors (Area III), each containing perturbations across a wide range of doses and durations. This pattern indicates that these groups are organized by MoAs rather than experimental conditions, consistent with the findings from LINCS [11]. Beyond these MoA-homogeneous clusters, DEPICT also reveals finer substructures. In Area IV, perturbations of RTK inhibitors and PI3K–AKT–mTOR pathway inhibitors form a mixed cluster. This is biologically consistent with RTKs acting upstream of the PI3K–AKT–mTOR pathway, but it also suggests that, in DEPICT’s prediction, some RTK inhibitors are transcriptionally closer to PI3K–AKT–mTOR inhibitors than to other RTK compounds. Such cross-MoA clusters may reflect shared downstream signaling or off-target activities that are not apparent from MoA alone.

**Fig. 3:**
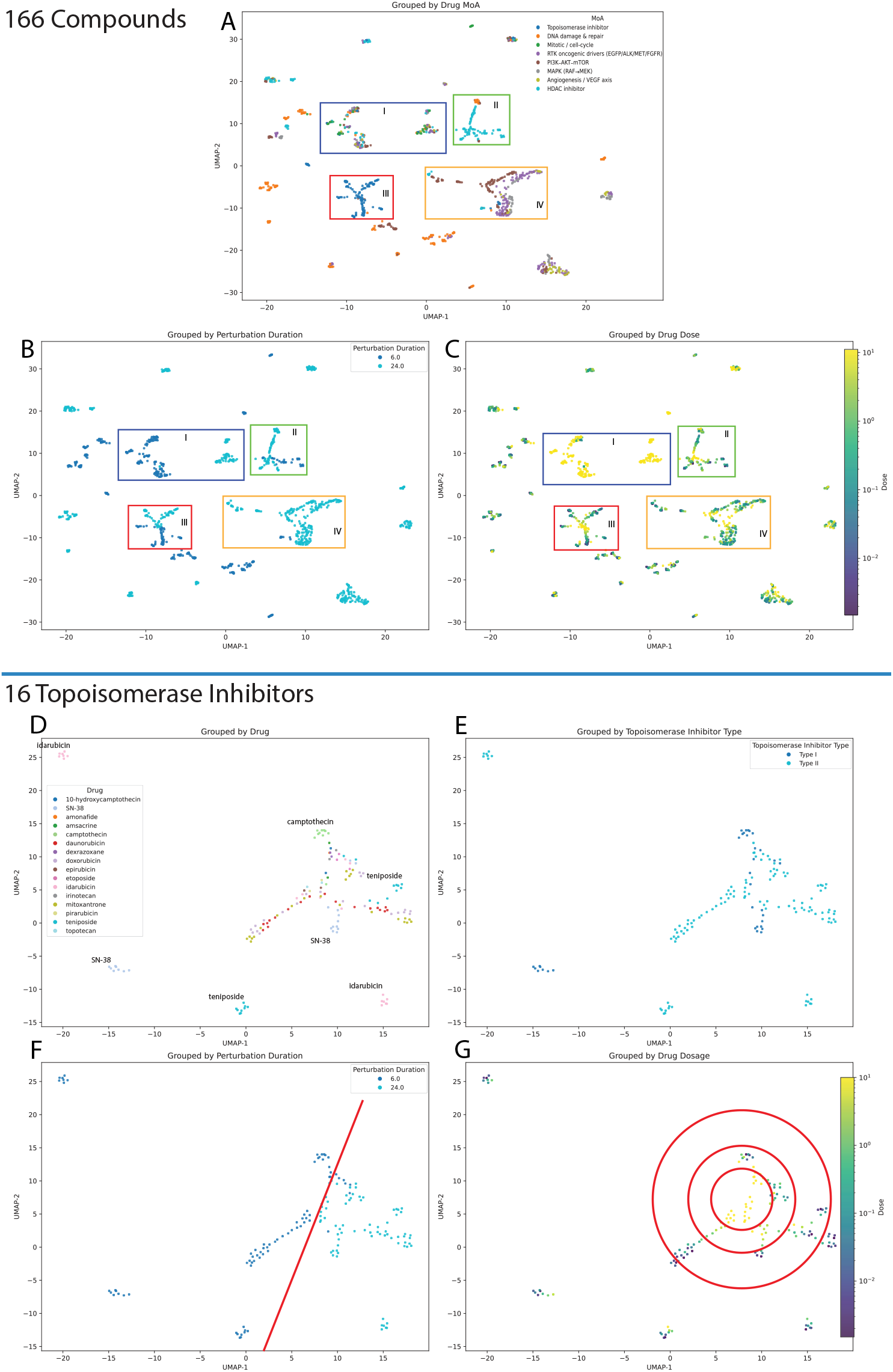
2D plot of predicted transcriptional responses on A549 11

As we found the topoisomerase inhibitors form a distinct cluster and topoisomerase inhibitors have also been widely used as treatment for various cancers [57], we further focused on the analysis of 150 perturbations from 16 topoisomerase inhibitors. In addition to forming a distinct cluster at the global level, topoisomerase inhibitors exhibit clear substructure. Several compounds (Idarubicin, SN-38, Teniposide, Camptothecin) separate from the main topoisomerase cloud (Fig. 3D). This indicates that DEPICT captures coarse subtype-specific transcriptional variation within a single MoA class, not only the broad distinction between different MoAs. In Fig. 3E, a limited but suggestive separation between type I and type II inhibitors supports this, although the evidence is constrained by the small number of type I compounds.

Condition effects are also well indicated in DEPICT. In Area I of the global plot (Fig. 3B), perturbations with similar MoAs and high concentrations split into two adjacent duration-driven subclusters, with 6-hour treatments on one side and 24-hour treatments on the other. A similar pattern appears for perturbations annotated as “DNA damage & repair,” which form multiple small clusters where duration is the same but doses vary. Within the topoisomerase subset, duration and dose gradients are clearer. In Fig. 3F, predicted responses are approximately separated by duration (6 vs. 24 hours), while in Fig. 3G, dose organizes the plot in a radial manner, with higher concentrations enriched near the center, and lower concentrations pushed to the outer rings.

This exploratory analysis also suggests several preclinical hypotheses for followup experiments. First, it can be used to explore sub-MoA heterogeneity within topoisomerase inhibitors. Comparing the isolated drugs with those in the main topoisomerase cluster could reveal whether these compounds truly engage distinct biological programs or primarily reflect off-target effects. Second, the mixed RTK–PI3K–AKT–mTOR cluster in Area IV harbored drugs with different MoAs but similar predicted transcriptional responses. If this enrichment is not driven by off-target activity, the RTK drugs in this region may be worth a detailed evaluation for drug repurposing or combination therapy with PI3K–AKT–mTOR inhibitors. In summary, this exploratory UMAP analysis shows that DEPICT not only reproduces known MoA-level organization but also reveals subtle subtype clusters and smooth gradients from dose and duration. These structures provide a map for designing follow-up experiments such as dissecting subtype-specific mechanisms and identifying off-target.

## Discussion

In this study, we developed DEPICT, a deep learning framework for predicting condition-matched transcriptional responses to pharmacological perturbations. Across multiple evaluation settings, DEPICT showed improved generalization to unseen drugs and cellular contexts and achieved strong predictive performance for drug-induced differential gene expression. Beyond benchmark performance, our findings suggest that condition-matched in silico perturbation profiling may help extend the utility of perturbational transcriptomics for translational oncology. By enabling systematic exploration of drug responses across biological contexts and exposure settings, DEPICT provides a scalable framework for early stage therapeutic discovery and hypothesis generation. This capability is particularly relevant because a persistent challenge in oncology drug discovery is the difficulty of evaluating therapeutic candidates across the combinatorial space defined by tumor context, compound identity, dose and duration. Although perturbational resources such as LINCS L1000 have enabled large scale study of drug-induced transcriptional responses, their coverage of possible drug–cell–condition combinations remains sparse. As a result, many biologically and clinically relevant settings are not directly measured, limiting the use of perturbational transcriptomics for translational tasks. This gap is difficult to close experimentally because systematic profiling across many drugs and exposure conditions is not only impractical because of cost, but also often ethically constrained in human subjects. By computationally generating condition-matched perturbation profiles, DEPICT helps address this bottleneck and expands the range of settings in which transcriptomic response hypotheses can be explored.

Our findings also highlight the potential role of artificial intelligence in supporting preclinical drug discovery. In the NSCLC case study, DEPICT-based virtual screening prioritized compounds predicted to reverse disease-associated transcriptional signatures, and most of the top-ranked candidates had prior clinical or preclinical support in NSCLC. These findings suggest that predictive models of perturbational transcriptomics may serve as practical tools for narrowing large chemical spaces to smaller sets of biologically plausible therapeutic hypotheses. More broadly, such models may complement experimental screening by prioritizing candidates before laboratory validation and helping focus resources on compounds with stronger disease relevance. In settings where exposure conditions are central, such as drug synergy modeling, DEPICT may also provide condition-matched perturbation profiles when experimentally measured signatures are unavailable.

Beyond predictive utility, the transcriptional landscapes generated by DEPICT may provide a useful view of the organization of drug-induced cellular responses. In our exploratory analyses, perturbations showed broad clustering by mechanism of action, suggesting that the model captures biologically meaningful relationships among compounds. Mixed clusters involving drugs targeting related signaling pathways may reflect shared downstream transcriptional programs or other mechanistic connections that warrant further study. These observations indicate that predicted perturbation profiles may be useful not only for ranking candidate therapies but also for generating hypotheses about drug mechanisms, pathway-level interactions, and potential combination strategies.

Several limitations should be considered when interpreting these findings. First, DEPICT was trained on transcriptional responses measured primarily in cancer cell lines. Although cell-line systems provide a scalable and controlled platform for perturbation modeling, they do not fully capture the complexity of human tumors, including microenvironmental influences, immune interactions, and inter-patient heterogeneity [58, 59]. Second, the model is based on bulk microarray-like LINCS L1000 profiles, which offer limited cellular resolution relative to single-cell measurements [60]. Third, although DEPICT improved predictive performance over existing approaches, its predictions remain imperfect and should not be interpreted as substitutes for experimental validation. Accordingly, DEPICT is best viewed as a tool for generating large scale, condition-matched transcriptional profiles that support translational applications such as preclinical prioritization and hypothesis generation, rather than direct clinical decision-making. Future work incorporating patient-derived models, organoids, single-cell perturbation datasets, and richer biological priors may further improve the translational relevance of predicted perturbational profiles.

Overall, our study shows that accurate prediction of condition-matched transcriptional responses is feasible and can support translationally relevant downstream analyses in oncology. As perturbational datasets continue to expand and predictive models improve, AI-assisted perturbation modeling may become an increasingly valuable complement to experimental screening for drug repurposing, combination discovery, and mechanism-guided therapeutic development.

## Methods

### Transcriptional data preprocessing

We used the LINCS L1000 Phase I database (GSE92742) [11], a publicly available large collection of transcriptional profiles spanning thousands of perturbagens across various durations, doses, and cell lines. LINCS L1000 measures bulk gene expression with a reduced-representation, ligation-mediated assay in which barcoded amplicons hybridize to color-coded Luminex beads to quantify 978 landmark genes. The remainder of the transcriptome is computationally inferred. The LINCS project used 384-well plates for the experiments. Each plate contained a single cell line, with several control wells treated with DMSO and the remaining wells treated with different perturbations. In this study, we restricted all analyses to the 978 landmark genes that were experimentally measured.

We preprocessed the dataset from the level-3 LINCS dataset used in PRnet [16]. PRnet excluded perturbations with insufficient numbers of compound (obsevations *<* 5) and removed perturbations with invalid SMILES. Based on the PRnet dataset, we created two datasets. First, we constructed a small dataset to facilitate model tuning with reduced computational cost. From the cleaned PRnet dataset, we further filtered and kept the top 10 cell lines with the most perturbations, drugs can be traced in PubChem, and perturbations under the condition of 10 *µ*M and 24 hours. This small dataset consisted of 85,576 perturbations and 25,661 controls from 1,479 plates, and contained 10 cell lines, 2,590 drugs and one perturbation condition (10-*µ*M and 24-hour). Second, we constructed a full dataset to train the final DEPICT model. From the cleaned PRnet dataset, we further excluded plates without any control wells. The resulting full dataset comprised 836,649 perturbations and 46,428 controls from 2,875 plates, covering 82 cell lines, 17,203 drugs, and various durations and dose levels. We then constructed baseline–perturbation pairs for both the small and full dataset. For each perturbation profile, we randomly selected one control profile from the same plate to serve as its baseline, yielding one matched pair per perturbation profile.

### Drug representations

Canonical SMILES strings for each compound were obtained from the LINCS L1000 metadata. Mechanism of action (MoA), targets, clinical indications, and other biomedical annotations were retrieved from the Drug Repurposing Hub [61]. We represented each compound using two complementary embeddings, which are 512-bit binary Morgan fingerprint and 512-dimensional LLM-derived continuous embedding. Morgan fingerprints were computed from canonical SMILES strings using RDKit [62]. To generate the LLM-derived embeddings, we first constructed a compact augmented text description for each compound, including its name, PubChem CID, SMILES, mechanism of action, targets, and clinical context, using a large language model (“gpt-4o”). We then encoded this text using the embedding model “text-embedding-3-large” to obtain a 512-dimensional continuous vector for each compound. Details of the LLM-based embedding procedure, including the prompt, are provided in Code Availability.

### Model architecture

The DEPICT framework is illustrated in Fig 1, Phase II. A more detailed technical description of the DEPICT architecture and its constituent modules is provided in Appendix A, Fig A1. DEPICT comprises three encoder components and a prediction head. First, we compute the mean (*µ*_*i*_ ∈ ℝ^978^) and variance (*V*_*i*_ ∈ ℝ^978^) of baseline expression for each gene within a given cell line. The baseline expression profile for the *i*-th sample ((*X*_*i*_, *µ*_*i*_, *V*_*i*_) ∈ ℝ^978^× ℝ^3^) is then passed to a gene-specific encoder (*f*_*gene*_ : ℝ^978^ × ℝ^3^ →ℝ^978^ × ℝ^32^). The gene-specific encoder consists of 978 multilayer perceptrons (MLPs) with identical architectures, with each MLP fitted separately for every single gene. This gene-specific encoder allows DEPICT to learn richer latent representations of gene expression profiles 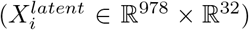 at the single-gene level rather than modeling all genes jointly. A vanilla transformer encoder (*f*_*tf*_ : ℝ^978^ × ℝ^32^ → ℝ^978^ × ℝ^32^) is then used to model the interactions among genes, generating latent gene features that incorporate gene-gene relationships 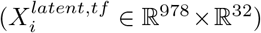.

In parallel, Morgan fingerprints (*D*_*MF*,*i*_ ∈ {0, 1}^512^) and LLM embeddings (*D*_*LLM*,*i*_ ∈ ℝ^512^) are processed by separate encoders with the same architecture (*f*_*mf*_ : {0, 1}^512^ → ℝ^128^ and *f*_*llm*_ : ℝ^512^ → ℝ^128^), producing refined low-dimensional latent drug features 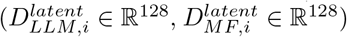. These latent gene and drug features are subsequently integrated into a gene-drug fusion encoder (*f*_*fuse*_ : ℝ^978^ × ℝ^32^ × ℝ^128^ → ℝ^978^ ×ℝ^32^) to learn latent perturbed gene features 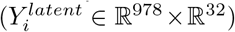, conditioned on the dose (*c*_*i*_) and duration (*t*_*i*_) of the perturbation. Finally, the latent perturbed gene features are passed to the prediction head (*f*_*pred*_ : ℝ^978^ × ℝ^32^ → ℝ^978^) to predict the perturbed gene expression profiles (Ŷ_*i*_ ℝ^978^). Additional details on the model architecture and tuned hyperparameters are provided in Appendix A.

### Gene-specific encoder and transformer encoder

The gene-specific encoder maps each gene’s profile ((*X*_*i*_, *µ*_*i*_, *V*_*i*_) ∈ ℝ^978^ × ℝ^3^) into a richer latent space 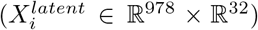. For the profile of the *j*-th gene in the *i*-th perturbation ((*X*_*i*,*j*_, *µ*_*i*,*j*_, *V*_*i*,*j*_) ∈ ℝ^3^), a distinct two-layer feed-forward network (*f*_*gene*,*j*_ (*X*_*i*,*j*_, *µ*_*i*,*j*_, *V*_*i*,*j*_) : ℝ^3^ → ℝ^32^) is fitted to generate a richer latent feature for the *j*-th gene 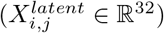. Thus, the gene-specific encoder comprises 978 separate small MLPs, one for each gene. Concatenating these features across genes yields the latent feature tensor for the *i*-th perturbation 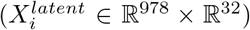. This encoder enables DEPICT to learn fine-grained features at the level of individual genes rather than learning a coarse space that embeds all genes.

The enhanced gene-specific latent features 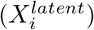 are then passed through four stacked vanilla transformer encoder layers (*f*_*tf*_ () : ℝ^978^ × ℝ^32^ → ℝ ^978^ × ℝ^32^) [14]. The relationships among genes are modeled through self-attention, while the shared gene-level patterns are captured by the shared feed-forward sublayers in the transformer encoder 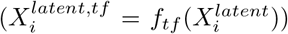 . A key limitation of the LINCS L1000 setting is that only 978 genes are directly measured, rather than the full transcriptome, which makes prior knowledge of gene relationships, such as pathway information, incomplete or noisy. Unlike GNN-based encoders [17, 18, 20], self-attention allows DEPICT to model gene interactions in a data-driven manner without being affected by lack of prior knowledge.

### Drug feature encoder

The drug feature encoder is a three-layer feed-forward network. Two separate encoders with the same architecture are used to process the binary 512-bit Morgan fingerprints (*D*_*MF*,*i*_ 0, 1 ^512^) and the continuous 512-dimensional LLM embeddings (*D*_*LLM*,*i*_ *∈* ℝ^512^), respectively. These encoders produce dense low-dimensional latent drug features 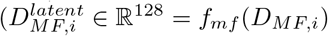 and 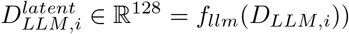.

### Gene-drug fusion encoder

The gene-drug fusion encoder comprises two stacked layers. Although the two layers are distinct, they share the same architecture. Each layer contains a multi-head cross-attention block and a scalar gating signal generated by a small MLP that takes as input the logarithms of dose and duration, log_10_(*c*_*i*_) and log_10_(*t*_*i*_), for the *i*-th perturbation. Specifically, the latent drug features (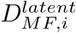 and 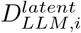) modulate the latent features of each gene 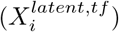 through cross-attention, thereby generating latent perturbed gene features. The gating signal then scales the output of the cross-attention block, producing latent perturbed gene features conditioned on the dose and duration of the perturbation 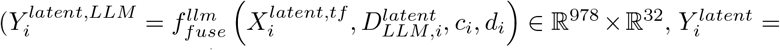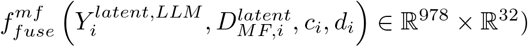.

### Prediction head

After generating the latent perturbed gene features, the prediction head maps each gene’s perturbed features to a scalar output using a gene-shared feed-forward network, a sequence of feature-modulation (FiLM) layers [63] that consider cell- and gene-specific adaptations, and two parallel per-gene readout paths (linear and residual MLP). For the *i*-th perturbation and *j*-th gene, the latent perturbed feature 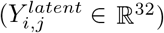 is first passed through a shared feed-forward network 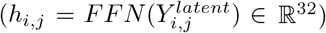.

To adapt predictions according to cell-specific uncertainty, the hidden representation (*h*_*i*,*j*_) is then modulated by the per-gene, cell-specific variance (*V*_*i*,*j*_) through the first FiLM layer.

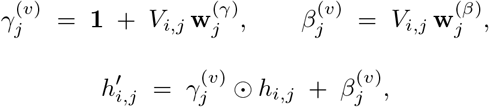

where 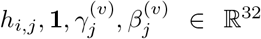, and ⊙ denotes element-wise multiplication. Here 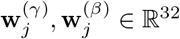 are learnable per-gene vectors.

A second FiLM layer introduces additional gene-specific adaptation through learnable scale and shift parameters:

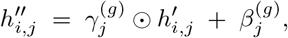

where 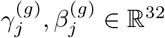 are the trainable parameters per gene.

The modulated hidden state 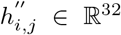 is then mapped to scalar predictions through two complementary paths: a linear path and a residual MLP path. The linear path projects each gene embedding to a scalar through a per-gene linear projection:

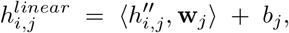

where **w**_*j*_ ∈ ℝ^32^, *b*_*j*_ ∈ ℝ are trainable parameters, and ⟨·, ·⟩ denotes the inner product.

In parallel, a two-layer MLP, parameterized separately for each gene, produces an additional scalar output:

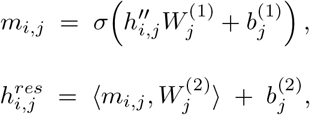

where 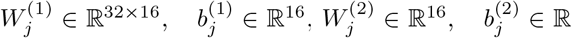 are trainable parameters, and *σ*(·) is the GELU (Gaussian Error Linear Unit) activation function.

Finally, the outputs of the two paths are summed to produce the predicted perturbed gene expression for each gene:

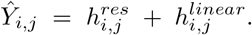

Concatenating predictions across genes yields the predicted perturbed expression profile for the *i*-th perturbation (Ŷ_*i*_ ∈ ℝ^978^).

### Training and testing

We minimize a composite objective that reduces the mean squared error (MSE) per gene and increases the Pearson correlation coefficient (PCC) between observed and predicted differential expression at the per-perturbation level:

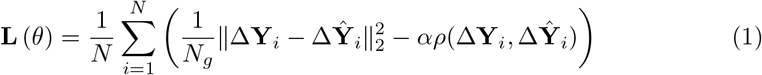

where Δ**Y**_*i*_ and Δ**Ŷ** _*i*_ ∈ ℝ^978^ denote the vectors of observed and predicted differential gene expression, respectively, for the *i*-th perturbation profile in the training set; *θ* denotes all trainable parameters in DEPICT; *N* and *N*_*g*_ denote the numbers of perturbation profiles and genes, respectively; *α* is a hyperparameter; and *ρ*(·) denotes the Pearson correlation.

This loss function encourages accurate prediction while preserving the directionality of gene-level changes. Additional hyperparameter settings such as *α*, learning rate, batch size, and so on are provided in Appendix A. Before training, we normalized gene expression within each perturbation profile. Specifically, each profile was divided by the sum of its gene expression values across all genes, so that all profiles had the same total expression after normalization.

The small dataset described in Transcriptional data preprocessing was used for large-scale hyperparameter tuning of DEPICT and to identify a reduced hyperparameter grid. DEPICT was then trained and further tuned on the full dataset using this smaller grid.

All datasets were split into training, validation, and test sets in an 8:1:1 ratio under three strategies, which were random split (by perturbation), drug split (by compound), and cell split (by cell line). In drug or cell splits, compounds or cell lines appearing in one set were excluded from the other two sets. Examples of different split strategies are illustrated in Appendix A Fig A2. These data-splitting strategies were designed to evaluate how the model generalizes to unseen perturbations, compounds, and cell lines. For each strategy, we used five-iteration Monte Carlo cross validation, training all models under five independent data splits to obtain robust comparative results. Within each repetition, all models were trained and evaluated on identical datasets. Hyperparameters were tuned based on the performance in the validation set, whereas the test set was used only for the final model performance comparison and downstream analyses.

### Model evaluation

We assessed the predictive performance of each model using three metrics: per-gene mean squared error (MSE), expected Pearson correlation coefficient (PCC) per sample, and expected coefficient of determination (*R*^2^) per sample.

1. MSE per gene, *MSE*_*g*_(**Y**_*i*_, **Ŷ**_*i*_):

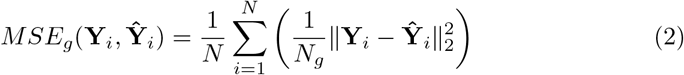
2. Expected PCC per sample, *PCC*(**Y**_*i*_, **Ŷ**_*i*_):

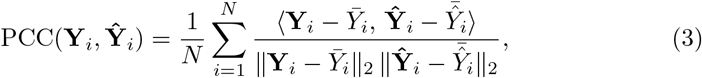
3. Expected *R*^2^ per sample, *R*^2^(**Y**_*i*_, **Ŷ**_*i*_):

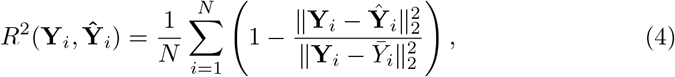

where ∥· ∥ denotes the Euclidean norm, ⟨·, ·⟩ denotes the inner product; **Y**_*i*_ denotes the observed perturbed gene expression; 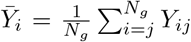 and 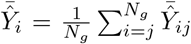 denote the mean observed and predicted gene expression for a profile vector, respectively.

Note that *MSE*_*g*_(**Y**_*i*_, **Ŷ** _*i*_) = *MSE*_*g*_(Δ**Y**_*i*_, Δ**Ŷ** _*i*_), but this equivalence does not hold for *PCC*(·,·) and *R*^2^(·, ·).

Finally, we used *MSE*_*g*_(**Y**_*i*_, **Ŷ** _*i*_), *PCC*(**Y**_*i*_, **Ŷ** _*i*_), *PCC*(Δ**Y**_*i*_, Δ**Ŷ** _*i*_), *R*^2^(**Y**_*i*_, **Ŷ** _*i*_), and *R*^2^(Δ**Y**_*i*_, Δ**Ŷ** _*i*_) as evaluation metrics. For simplicity, these metrics are denoted as MSE, PCC, ΔPCC, *R*^2^ and Δ*R*^2^ in the benchmarking performance Table 1.

### Simple baselines

Ahlmann-Eltze et al. [23] recently showed that sophisticated learning models can be outperformed by simple baselines. For example, when predicting transcriptional responses to genetic perturbations, directly using baseline gene expression as the predicted perturbed expression can outperform state-of-the-art deep learning methods.

Motivated by this observation, we included several simple but informative baselines to benchmark the performance of deep learning models.

The first baseline, denoted “Naive”, uses the baseline (unperturbed) gene expression profile as the predicted perturbed expression for each experiment in the test set. The second baseline, termed “Train Mean (TM)”, uses the average perturbed gene expression profile computed over all perturbations in the training set as the prediction for every test perturbation. We also defined three conditional mean baselines as “Train Mean across Cell (TM-C)”, “Train Mean across Drug (TM-D)”, and “Train Mean across Cell-Drug Pair (TM-CD)”. TM-C predicts the perturbed expression for a test perturbation as the mean perturbed expression over all training perturbations with the same cell line. TM-D analogously uses the mean over all training perturbations with the same drug, and TM-CD uses the mean over all training perturbations with the same cell–drug pair.

### Mechanism-guided virtual screening identifies candidates that reverse NSCLC transcriptional signatures

We evaluated whether DEPICT-predicted transcriptional responses in the LINCS L1000 framework could support in silico screening of candidate compounds for single-agent therapy. Specifically, we tested whether a compound’s predicted effect in the non-small cell lung cancer (NSCLC) cell line A549 could reverse the NSCLC disease signature to a more normal-like state.

The NSCLC disease signature was derived from a published dataset [64], accessed via Expression Atlas [65]. The reference dataset reports genome-wide tumor-versus-normal log-fold changes from paired NSCLC samples. We intersected these genes with the LINCS L1000 landmark gene set and retained only those with statistically significant log-fold changes in the reference dataset, yielding 690 genes. The resulting NSCLC signature was then a 690-dimensional vector of tumor-versus-normal log-fold changes representing the disease state. We then constructed a “reversal” signature by inverting the direction of deregulation (equation 5). Intuitively, genes up-regulated in tumors should be down-regulated, and vice versa. This reversed NSCLC signature served as the reference for subsequent scoring of predicted drug responses.

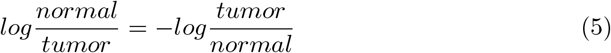

We predicted differential expression profiles in A549 for all 17,203 compounds. For each compound, we predicted its profiles under three commonly assayed conditions, which are 10 *µ*M for 6 h, 10 *µ*M for 24 h, and 5 *µ*M for 24 h. For the downstream analysis, each profile was restricted to the 690 genes included in the NSCLC reversal signature. We then scored every compound’s transcriptional profiles and took the best rank across the three conditions for each compound.

To prioritize candidate compounds, we quantified the alignment between each predicted compound profile and the reversal signature using two rank-based similarity metrics. Rank-based metrics help mitigate the differences in scale and measurement platform between LINCS and the reference dataset. The first metric was the Spearman rank correlation between a compound’s predicted differential expression profile and the NSCLC reversal signature. Higher value indicates better directional consistency of a compound’s ability to reverse the disease state. The second metric was the GSEA-style connectivity score [66]. From the reversal signature, we defined two disjoint gene sets: 285 genes that should be up-regulated and 405 genes that should be down-regulated to oppose the disease state. For each compound profile, we ranked the 690 genes by predicted differential expression and computed enrichment scores to assess whether genes targeted for up-regulation were enriched at the top of the ranking, while those targeted for down-regulation were concentrated at the bottom. Then these two enrichment scores were then combined into a single connectivity score. Higher connectivity scores indicate that the predicted transcriptional changes induced by a compound align more closely with the changes needed to reverse the NSCLC transcriptional profile.

1. Spearman correlation between the reversed disease signature and each drug profile, SPC(Δ**Ŷ** _*i*_, **S**):

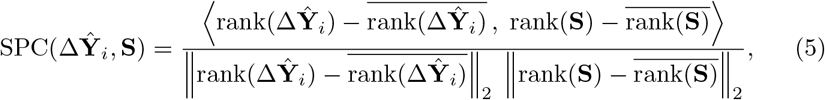

where Δ**Ŷ** _*i*_ ∈ ℝ^690^ is the predicted differential expression for *i*-th profile, and **S** ∈ R^690^ is the reversed NSCLC signature as reference. ra<u>nk(·) re</u>turns the rank of each gene within the given vector (average ranks for ties), rank(·) is the mean rank broadcast across the vector, and ⟨·,·⟩ is the inner product.
2. GSEA-style connectivity score [66] between the reversed disease signature and each drug profile, Conn_*i*_:

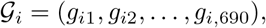

where *g*_*ij*_ is the *j*-th gene when all 690 genes are sorted in decreasing order of Δ**Ŷ** _*i*_.

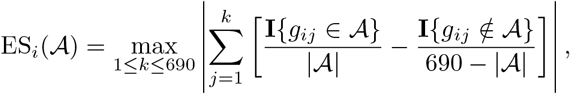

where ES_*i*_(𝒜) is the calculated enrichment score for the *i*-th profile, **I**{·} is the indicator function, and 𝒜 ⊆ 𝒢 _*i*_ is a predefined gene set. For the reversed disease signature, let 𝒰 be the 285 genes requiring up-regulation and 𝒟 be the 405 genes requiring down-regulation. We then define the connectivity score for the *i*-th profile as:

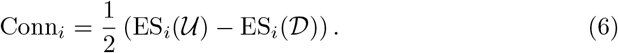
3. Finally, we calculate the ranking score *RS*_*i*_ for the *i*-th profile as the sum of the Spearman correlation and the connectivity score:

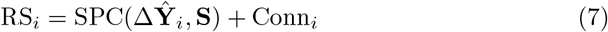

For the final ranking step, we used the ranking score *RS*_*i*_, where a higher values indicate a stronger predicted reversal of the transcriptional response associated with NSCLC. As a benchmark, we applied the same scoring procedure to the observed LINCS differential expression profiles of the cell line A549, and compared the rankings of the observed profiles with the rankings of the DEPICT-predicted profiles.

### Synergy prediction using condition-matched predicted profiles

The reference dataset used to label drug-doublet synergy status was obtained from O’Neil et al. [54]. In this dataset, each single drug and drug doublet was evaluated under multiple dose regimens and 96-hour duration in the HT29 cell line. After intersecting compounds between the reference dataset and LINCS L1000, we identified 19 overlapping drugs and 556 unique pairs that have perturbations in HT29. These 556 pairs were then annotated as synergistic or antagonistic according to their Loewe additivity scores [67], calculated using the Python package “synergy” [68].

To construct predictive features, we used the DEPICT-predicted transcriptional response for each drug under the dose regimen and duration specified in the reference dataset. Each response vector comprised 978 gene-level features. Concatenating the two single-drug vectors yielded a 1,956-dimensional representation for each pair, which was high relative to the sample size. To reduce dimensionality and mitigate overfitting, we applied PCA [69] to project each single-drug profile to 50 dimensions, producing a 100-dimensional concatenated input feature for each drug doublet, paired with a binary outcome (synergistic or antagonistic).

We evaluated DEPICT-predicted transcriptional responses using two classical classifiers, which are ridge logistic regression [70] and random forest [71]. Performance was assessed using leave-one-out cross-validation with area under the receiver operating characteristic curve (AUC), area under the precision-recall curve (PR-AUC), accuracy and macro-F1 as evaluation metrics. For comparison, we repeated the same classification task using experimentally measured LINCS L1000 transcriptional profiles. Because many specific dose and duration combinations required by the reference dataset, were unavailable for HT29 in the observed LINCS data, we used the closest available LINCS conditions as a proxy when exact matches were not available.

### Exploratory analysis on predicted perturbations

The MoA annotations were obtained from the Drug Repurposing Hub [61]. Focusing on NSCLC, we selected the MoAs commonly implicated in NSCLC biology, including receptor tyrosine kinase (RTK) inhibitors, microtubule inhibitors, angiogenesis/VEGF inhibitors, MAPK pathway (RAF→ MEK) inhibitors, PI3K-AKT-mTOR inhibitors, histone deacetylase (HDAC) inhibitors, topoisomerase inhibitors, and agents targeting DNA damage and repair. A total of 166 drugs with above MoAs were selected. We then screened these drugs in the A549 cell line under perturbation conditions (dose and duration) available in the observed LINCS L1000 dataset. For each perturbation, the baseline expression was computed as the average across all DMSO controls run on the same plates under matched conditions. DEPICT was then used to predict the differential gene expression (DGE) for each perturbation.

The predicted DGEs were visualized in two-dimensional space using principal component analysis (PCA) [69] and uniform manifold approximation and projection (UMAP) [72]. First, the 978-gene expression space was reduced to 50 principal components using PCA. In this PCA space, a k-nearest neighbor (kNN) [73] graph was constructed using the cosine distance, and then a two-dimensional UMAP embedding was computed from the precomputed neighbor graph. The resulting two-dimensional coordinates were plotted as scatter points and annotated with perturbation attributes, including compound, MoA, perturbational duration, and dose.

## Declarations

### Data Availability

The original LINCS L1000 (GSE92742) profiles can be downloaded from https://www.ncbi.nlm.nih.gov/geo/query/acc.cgi?acc=GSE92742.

The partial LINCS L1000 dataset used in this study was downloaded from PRNet preprocessed data: https://zenodo.org/records/14230870.

The metadata for drugs were retrieved via Drug Repurposing Hub: https://repo-hub.broadinstitute.org/repurposing#download-data.

The preprocessed data are available on GitHub at https://github.com/lazypuff/DEPICT and on Zenodo at https://zenodo.org/records/19207077.

### Code Availability

Codes implementing DEPICT algorithm are publicly available in Github: https://github.com/lazypuff/DEPICT.

## Appendix A Detailed model structure

Fig A1 is the comprehensive architecture of the DEPICT framework. The detailed explanation for every notation used in this paper is listed in Table A1 and A2.

**Table A1:** Notations in DEPICT.

| Symbol | Space | Definition |
| --- | --- | --- |
| **Input gene features** |  |  |
| $X_i$ | $\mathbb{R}^{978}$ | Baseline expression profile for perturbation $i$. |
| $\mu_i$ | $\mathbb{R}^{978}$ | Cell-specific mean expression for perturbation $i$. |
| $V_i$ | $\mathbb{R}^{978}$ | Cell-specific variance of gene expression in perturbation $i$. |
| $(X_{i,j}, \mu_{i,j}, V_{i,j})$ | $\mathbb{R}^3$ | Concatenation of scalar inputs for gene $j$ in perturbation $i$. |
| **Gene latent representations** |  |  |
| $X_i^{latent}$ | $\mathbb{R}^{978} \times \mathbb{R}^{32}$ | Per-gene latent features from gene-specific encoders for perturbation $i$. |
| $X_i^{latent,tf}$ | $\mathbb{R}^{978} \times \mathbb{R}^{32}$ | Transformer-enhanced gene latent features for perturbation $i$. |
| **Drug features** |  |  |
| $D_{MF,i}$ | $\{0, 1\}^{512}$ | Morgan fingerprint for the drug in perturbation $i$. |
| $D_{LLM,i}$ | $\mathbb{R}^{512}$ | LLM-derived embedding for the drug in perturbation $i$. |
| $D_{MF,i}^{latent}, D_{LLM,i}^{latent}$ | $\mathbb{R}^{128}$ | Encoded low-dimensional drug features for perturbation $i$. |
| **Perturbation conditions** |  |  |
| $c_i$ | $\mathbb{R}$ | Compound dose for perturbation $i$. |
| $t_i$ | $\mathbb{R}$ | Duration for perturbation $i$. |
| **Fusion and prediction representations** |  |  |
| $Y_i^{latent}$ | $\mathbb{R}^{978} \times \mathbb{R}^{32}$ | Latent perturbed gene features after fusion for perturbation $i$. |
| $h_{i,j}$ | $\mathbb{R}^{32}$ | Shared-MLP transformed representation for perturbation $i$, gene $j$. |
| $h'_{i,j}, h''_{i,j}$ | $\mathbb{R}^{32}$ | FiLM-modulated hidden representations for perturbation $i$, gene $j$. |
| **FiLM parameters** |  |  |
| $\gamma_j^{(v)}, \beta_j^{(v)}$ | $\mathbb{R}^{32}$ | Variance-driven FiLM scale and shift for gene $j$. |
| $\gamma_j^{(g)}, \beta_j^{(g)}$ | $\mathbb{R}^{32}$ | Gene-specific FiLM scale and shift for gene $j$. |
| **Outputs** |  |  |
| $h_{i,j}^{linear}$ | $\mathbb{R}$ | Output from linear readout path for perturbation $i$, gene $j$. |
| $h_{i,j}^{res}$ | $\mathbb{R}$ | Output from residual MLP path for perturbation $i$, gene $j$. |
| $\hat{Y}_{i,j}$ | $\mathbb{R}$ | Predicted expression for for perturbation $i$, gene $j$. |
| $\hat{Y}_i$ | $\mathbb{R}^{978}$ | Predicted perturbed profile for perturbation $i$. |

**Table A2:** Notation for functions in DEPICT.

| Function | Mapping | Description |
| --- | --- | --- |
| $f_{\text{gene}}$ | $\mathbb{R}^{978} \times \mathbb{R}^3 \rightarrow \mathbb{R}^{978} \times \mathbb{R}^{32}$ | Gene-specific encoder composed of 978 per-gene MLPs. |
| $f_{\text{gene},j}$ | $\mathbb{R}^3 \rightarrow \mathbb{R}^{32}$ | Per-gene MLP encoding gene $j$. |
| $f_{\text{tf}}$ | $\mathbb{R}^{978} \times \mathbb{R}^{32} \rightarrow \mathbb{R}^{978} \times \mathbb{R}^{32}$ | Transformer encoder modeling gene-gene interactions. |
| $f_{mf}$ | $\{0, 1\}^{512} \rightarrow \mathbb{R}^{128}$ | Encoder for Morgan fingerprints. |
| $f_{llm}$ | $\mathbb{R}^{512} \rightarrow \mathbb{R}^{128}$ | Encoder for LLM-based drug embeddings. |
| $f_{\text{fuse}}$ | $\mathbb{R}^{978} \times \mathbb{R}^{32} \times \mathbb{R}^{128} \rightarrow \mathbb{R}^{978} \times \mathbb{R}^{32}$ | Gene-drug fusion encoder using cross-attention and dose-duration gating. |
| $FFN$ | $\mathbb{R}^{32} \rightarrow \mathbb{R}^{32}$ | Shared MLP applied to each gene before FiLM modulation. |

**Fig. A1:**
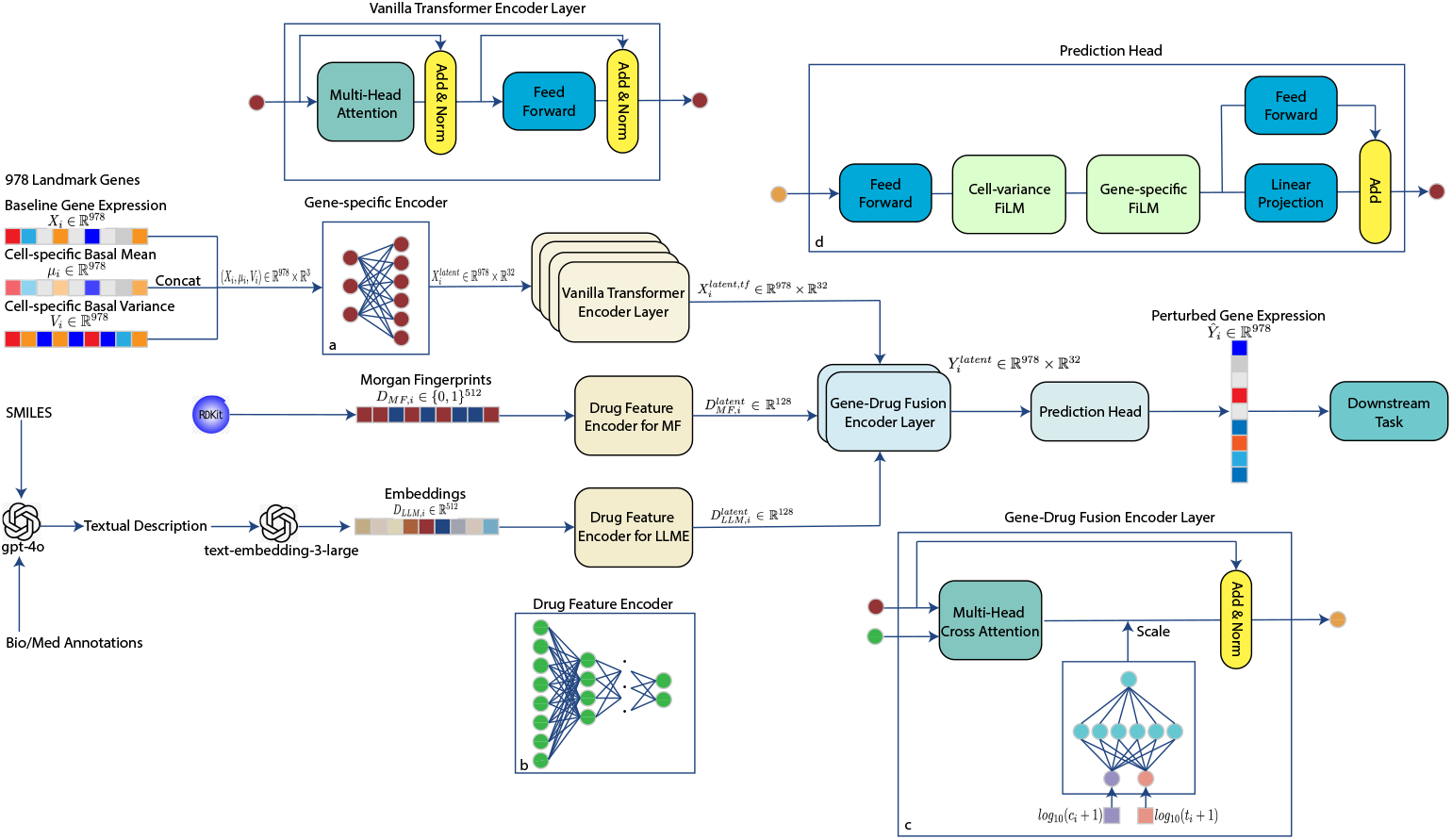
DEPICT architecture flowchart

Table A3 presents the detailed structural designs for each module in DEPICT, and Table A4 shows the tuning grid and selected values for the hyperparameters. In addition, DEPICT were trained using the AdamW optimizer and a cosine-annealing learning rate schedule with a warm-up start.

**Table A3:** Detailed model architecture and module-wise hyperparameters in DEPICT.

| Module | Architecture | Detailed hyperparameters |
| --- | --- | --- |
| Per-gene encoder | 2-layer MLP | $3 \rightarrow 16 + \text{GELU} \rightarrow 32$ |
| Transformer encoder layer | Vanilla transformer encoder layer [14] | Hidden size $d = 32$; 8 heads; FFN dim = 2048; dropout = 0.1 |
| Drug encoder | 3-layer MLP | $512 \rightarrow 256 + \text{GELU} + \text{BN} + 0.1\text{dropout} \rightarrow 128 + \text{GELU} + \text{BN} + 0.05\text{dropout} \rightarrow 128$ |
| Cross-attention layer | Q from gene encoder; K,V from drug encoder | Hidden size $d = 32$; 8 heads; dropout = 0.1 |
| Dose-duration layer | 2-layer MLP | $2 \rightarrow 16 + \text{GELU} \rightarrow 1 + \text{softplus}$ |
| Gene-shared FFN | 2-layer MLP | $32 \rightarrow 64 + \text{GELU} + 0.2\text{dropout} \rightarrow 32$ |
| FiLM layer | Feature-wise linear modulation [63] | Hidden dim $d = 32$ |
| Per-gene prediction head (linear path) | Linear projection | $32 \rightarrow 1$ |
| Per-gene prediction head (residual path) | 2-layer MLP | $32 \rightarrow 16 + \text{GELU} \rightarrow 1$ |

**Table A4:** Hyperparameter tuning grid and selected values in DEPICT.

| Hyperparameter | Chosen value | Tuning grid |
| --- | --- | --- |
| Hidden size ($d$) in Transformer | 32 | {32, 64, 128, 256} |
| Dropout rate | 0.1 | {0, 0.05, 0.1, 0.2} |
| Attention heads | 8 | {4, 8, 16} |
| Number of Transformer layers | 4 | {1, 2, 3, 4, 6, 8} |
| Transformer FFN hidden dimension | 2048 | {64, 128, 256, 512, 1024, 2048} |
| Gene-shared FFN hidden dimension | 64 | {64, 128, 256} |
| Loss function: MSE $-\alpha$ PCC | $\alpha=0.3$ | $\alpha \in \{0.15, 0.3, 0.6, 1, 2\}$ |
| Batch size | 128 | {64, 128} |
| Learning rate | $1 \times 10^{-4}$ | $\{10^{-5}, 10^{-4}, 2 \times 10^{-4}, 5 \times 10^{-4}, 10^{-3}, 2 \times 10^{-3}, 5 \times 10^{-3}\}$ |

MLP: Multilayer perceptron.
GELU: Gaussian error linear unit.
BN: Batch normalization layer.
FFN: Feed-forward network.
Q, K, V: Queries, keys, and values used in the attention mechanism.
Softplus: Softplus activation function.
Hidden size ( $d$ ): Dimensionality of the latent representation in the transformer encoder.

Fig A2 is an illustrative example to explain the three different splitting strategies used in this study.

**Fig. A2:**
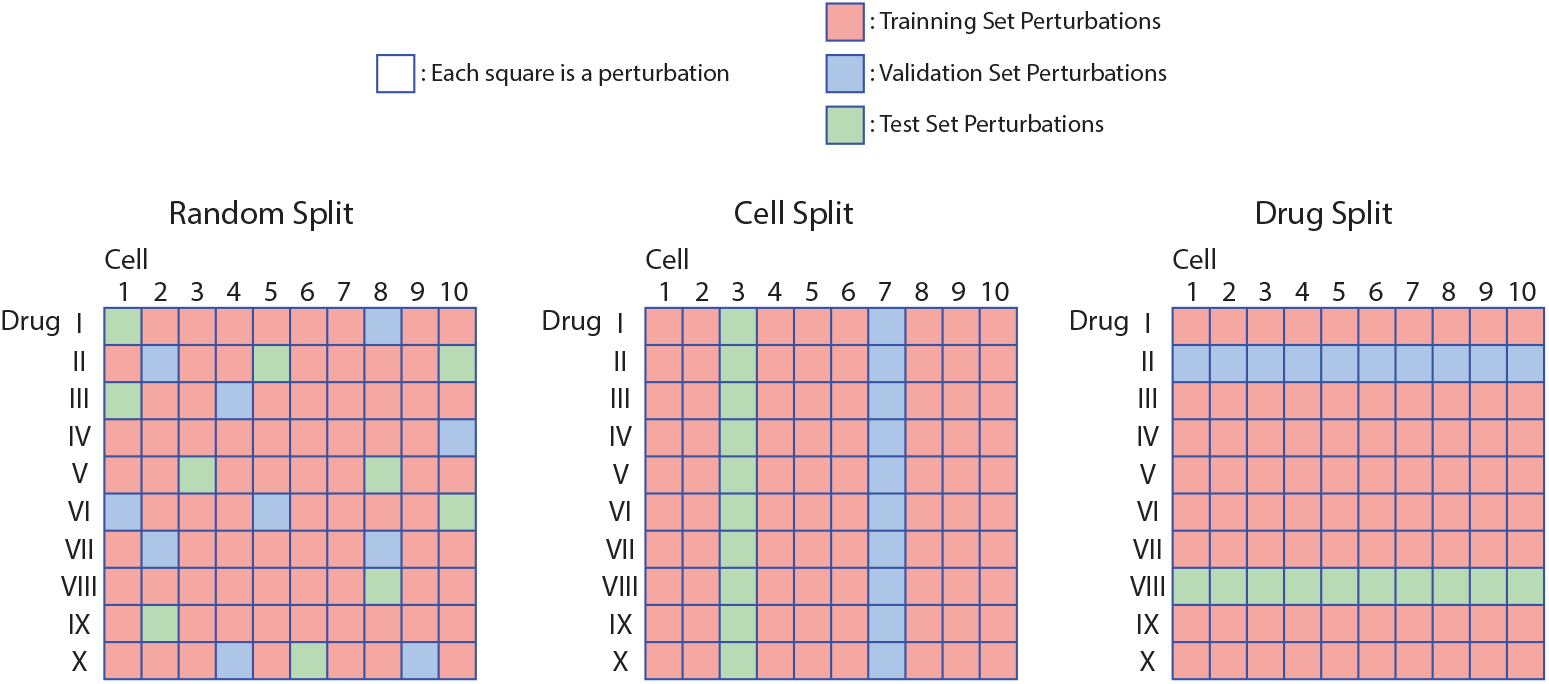
Types of split strategy

## APPENDIX B Detailed results

**Table B5:** List of Top-20 identified drugs and their mechanisms of action (MoA) for NSCLC cell line A549 based on predicted and observed LINCS L1000 data.

| Drug | MoA | Source |
| --- | --- | --- |
| AZD-8330 | MEK inhibitor | Predicted LINCS |
| MK-2206 | Akt inhibitor |  |
| PHA-793887 | CDK inhibitor |  |
| JW-7-24-1 | Not found |  |
| ZSTK-474 | Pan PI3K inhibitor |  |
| PI-828 | Pan PI3K inhibitor; CK2 inhibitor |  |
| foretinib | TKI |  |
| PI-103 | PI3K/Akt/mTOR inhibitor |  |
| dasatinib | DDR2/BRAF; TKI |  |
| KU-0060648 | Dual PI3K/DNA-PK inhibitor |  |
| midostaurin | TKI |  |
| JWE-035 | Not found |  |
| NVP-BEZ235/dactolisib | Dual PI3K/mTOR inhibitor |  |
| OSI-027 | mTOR inhibitor |  |
| VU-0418947-2 | Not found |  |
| CD-437 | Death receptor activator (DR4, DR5) |  |
| torin-2 | mTOR inhibitor |  |
| MRE-269 | Not found |  |
| GSK-2126458/omipalisib | Dual PI3K/mTOR inhibitor |  |
| BRD-K26381032 | Not found |  |
| wortmannin | PI3K inhibitor | Observed LINCS |
| NVP-BEZ235/dactolisib | Dual PI3K/mTOR inhibitor |  |
| NVP-TAE684 | ALK inhibitor |  |
| GDC-0941/pictilisib | Pan PI3K inhibitor |  |
| WYE-125132 | mTOR inhibitor |  |
| torin-2 | mTOR inhibitor |  |
| GSK-2126458/omipalisib | Dual PI3K/mTOR inhibitor |  |
| PI-103 | PI3K/Akt/mTOR inhibitor |  |
| AZD-7762 | Chk1 kinase inhibitor |  |
| epoxy-cholesterol | Not related to cancer |  |
| RITA | Activating p53 gene suppressor |  |
| AZD-8055 | mTOR inhibitor |  |
| dovitinib | FGFR inhibitor |  |
| mepacrine/quinacrine | p53/Wnt/PI3K/AKT/mTOR modulator |  |
| Crizotinib | ALK/ROS1 inhibitor |  |
| BMS-754807 | IGF-1R inhibitor |  |
| ZSTK-474 | Pan PI3K inhibitor |  |
| torin-1 | mTOR inhibitor |  |
| sirolimus | mTOR inhibitor |  |
| TGX-115 | PI3K inhibitor |  |

